# The synaptic vesicle protein Mover/TPRG1L is associated with lipid droplets in astrocytes

**DOI:** 10.1101/2022.01.18.476809

**Authors:** Jeremy Krohn, Florelle Domart, Thanh Thao Do, Thomas Dresbach

## Abstract

Crucial brain functions such as neurotransmission, myelination, and signaling pose a high demand for lipids. Lipid dysregulation is associated with neuroinflammation and neurodegeneration. Astrocytes protect neurons from lipid induced damage by accumulating and metabolizing toxic lipids in organelles called lipid droplets (LDs). LDs have long been considered as lipid storage compartments in adipocytes, but less is known about their biogenesis and composition in the brain. In particular, proteins covering the LD surface are not yet fully identified.

Here, we report that the synaptic vesicle protein Mover/TPRG1L, which regulates the probability of neurotransmitter release in neurons, is a component of the LD coat in astrocytes. Using conventional and super resolution microscopy, we demonstrate that Mover surrounds naive and oleic acid induced astrocytic LDs. We confirm the identity of astrocytic LDs using the neutral lipid stains Bodipy and LipidTox, as well as immunofluorescence for perilipin-2, a known component of the LD coat. In astrocytes, recombinant Mover was sufficient to induce an accumulation of LDs. Furthermore, we identified point mutations that abolish targeting to LDs and show similarities in the required binding sequences for association to the presynapse and LDs.

Our results show that Mover is not only a presynaptic protein but also a candidate for LD regulation. This highlights the dual role of Mover in synaptic transmission and regulation of astrocytic LDs, which may be particularly important in the context of lipid-related neurological disorders.

**Main points:**

- Mover is a lipid droplet protein
- Overexpression of Mover drives lipid droplet accumulation
- The F206 is important for LD targeting

## 1 Introduction

Mover (Mossy fiber associated vertebrate-specific protein; also called TPRG1L and SVAP30) is a non-transmembrane protein consisting of 266 amino acids expressed in different organs such as the heart, liver, and testis, with the highest expression being detected in the brain (Kremer *et al*. 2007; Burré *et al*. 2006). Mover is heterogeneously distributed in the mouse brain, with high abundance in the hippocampus, the amygdala, and the ventral pallidum (Wallrafen and Dresbach 2018). In neurons, Mover is located to the presynaptic terminal and associated to synaptic vesicles (Ahmed *et al*. 2013). Mover knockout mice show altered synaptic transmission at the calyx of Held and hippocampal mossy fiber terminals (Viotti and Dresbach 2019; Pofantis *et al*. 2021), implying that Mover modulates synaptic activity.

Synaptic transmission does not depend on neurons alone. Glial cells contribute to synapse formation and synaptic plasticity (Sancho *et al*. 2021), in part by forming a tripartite synapse, i.e. a connection between neuronal presynapse, postsynapse and astrocytes (Araque *et al*. 1999). Moreover, glial cells support neurons with metabolites, and recent studies demonstrate that they play a crucial role in various processes such as inflammation, elimination of waste products and neutralizing reactive oxygen species (ROS) (Giovannoni and Quintana, 2020; Islam *et al*. 2019). Astrocyte-neuron coupling is not limited to synaptic terminals. Cytotoxic lipid species that could cause damage to neuronal membranes and mitochondria are released via the apolipoprotein E (APOE) carrier and accumulate in astrocytes, where they can be stored in specialized lipid storage compartments called lipid droplets (LD) (Ioannou *et al*. 2019).

LD are lipid storage organelles found in nearly all species (Murphy 2012). They are highly abundant in adipose tissue, but are present in various other cell types as well. However, research of the past decade shed light on novel protective functions and implications for LDs in diseases (Welte 2015). Their evolutionary conservation renders LDs potential targets for the interaction between hosts and parasites such as Toxoplasma or Trypanosoma (Vallochi *et al*. 2018). Furthermore, the hepatitis-C virus harbors LDs for its own life cycle within the host cell (Miyanari *et al*. 2007). In zebrafish embryos, LDs are required for microbial defense before the immune system is fully active (Dutta *et al*. 2018). And research of the past decade shed light on novel protective functions and implications for LDs in diseases of the nervous system (Welte 2015). For example, in the Drosophila larval brain they protect a stem cell niche from hypoxia derived ROS damage (Bailey *et al*. 2015). Glia specific mitochondrial dysfunction in the adult Drosophila resulted in an accumulation of LDs and a progressive neurodegeneration (Cabirol-Pol *et al*. 2018), displaying LDs as a hallmark also for pathologic conditions, Ioannou *et al*. (2019) demonstrated that the accumulation of astrocytic LDs serves a neuroprotective function by sequestering toxic fatty acids derived from neurons. LDs metabolize lipids and form dynamic contacts with other cellular organelles for exchange of substrates (Valm *et al*. 2017).

LDs consist of a core, composed of triacylglycerols and sterol esters, that is surrounded by a phospholipid monolayer (Olzmann and Carvalho 2019). The monolayer surface is covered with proteins and enzymes that are necessary for trafficking, organelle interaction, lipid conversion, and regulation of LDs (Zehmer *et al*. 2009). Perilipins are a key group of LD proteins consisting of five isoforms, which regulate different aspects of LD generation and stabilization (Itabe *et al*. 2017). The protein composition of the LD surface is thought to contribute to the diversity of LD functions.

Here, we report that Mover is a novel component of the LD surface in astrocytes. Colocalization with LD markers demonstrated that endogenous Mover is indeed found surrounding oleic acid (OA) induced astrocytic LDs, and recombinant Mover retains the ability to target LDs. STED microscopy revealed a heterogeneous distribution around the LD surface with the formation of local clusters. We demonstrated that the overexpression of Mover in astrocytes induces an accumulation of LDs independent of OA induction. Furthermore, using different recombinant Mover constructs harboring point mutations, we identified necessary regions of Mover for LD binding. These results highlight that Mover not only modulates synaptic transmission at presynaptic terminals, but is also involved in LD regulation in astrocytes.

## 2 Materials and Methods

### 2.1 Plasmid constructs

All Mover constructs were designed using rat Mover sequences. The following constructs were used: Mover-5p-mut-EGFP, Mover-Y257P-EGFP, F206R-EGFP, Mover-4cam-EGFP, Mover-EGFP (promoter: CAG; backbone: pAM); Mover-DE2-mGFP, mGFP-52-266, mGFP-52-266-F206R, hsac2-mGFP, Mover-mGFP, Mover-myc, farnesylated GFP (GFP-F), mGFP, Mover-IRES-palmGFP (promoter: CMV; backbone: Clontech N1); mGFP-Mover (promoter: CMV; backbone: pCMV3).

### 2.2 Primary cell cultures

Primary hippocampal neurons and primary cortical astrocytes were obtained from 19 days old Wistar rat embryos as previously described by Dresbach *et al*. (2003) and Kaech and Banker (2006). Briefly, hippocampal or cortical cells were isolated in ice cold Hank’s Balanced Salt Solution (Gibco, HBSS) and dissociated by enzyme digestion with 0.05 – 0.25% Trypsin (Gibco) at 37°C for 20 min followed by mechanical dissociation. Hippocampal neurons were cultured at a density of 5 × 10^4^ cells.cm^−2^ on glass coverslips coated with 0.04% polyethyleneimine (Sigma, PEI) in Dulbecco⍰s Modified Eagle⍰s Medium (Gibco, DMEM) supplemented with 1% penicillin/streptomycin (Gibco) and 10% fetal bovine serum (Gibco, FBS), and the medium was changed to Neurobasal medium (Gibco, NB) supplemented with 1% penicillin/streptomycin, 0.0025% L-Glutamine (Gibco), and 2% B27 (Gibco) after one day. Cortical astrocytes were plated at a density of 2.5 × 10^4^ to 5 × 10^4^ cells.cm^−2^ in Minimum Essential Media (Gibco, MEM) supplemented with 10% Horse Serum (Gibco), 6 g/L Glucose (Sigma), and 200 mM L-Glutamine on glass coverslips coated with 1% PEI. The astrocyte culture medium was changed twice a week.

Human embryonic kidney cells (HEK293; RRID:CVCL_0063) were grown in 10 cm diameter culture dishes (Falcon) in DMEM supplemented with 1% penicillin/streptomycin and 10% FBS. Cells were passaged every two or three days. For immunofluorescence, cells were counted and plated at a density of 1-2 × 10^4^ in a 24-well plate on 0.04% PEI coated glass coverslips.

### 2.3 Transfection

HEK cells were transfected with several Mover constructs one day after plating using PEI transfection in Opti-MEM medium (Gibco). Per coverslip, 1 μg DNA was mixed with *1*% PEI in a 1:3 (v:v) ratio and incubated for 30 minutes at room temperature and carefully applied to each coverslip. Astrocyte cultures were transfected at day *in vitro* 13 using lipofectamine 2000 (Invitrogen) transfection following manufacturer’s instructions. The medium of the cells was changed with fresh pre-warmed medium before the transfection to ensure an exact volume per well. 0.8 μg DNA diluted in transfection buffer was incubated per coverslip. The medium was changed after 4 hours of incubation.

### 2.4 Oleic acid treatment

OA treatment was used to increase the number and size of LDs. OA complexes with bovine serum albumin (Sigma, BSA) were prepared with an adapted protocol from Listenberger *et al*. (2016). Briefly, a ~4.6 mM solution of OA-BSA complexes was obtained by adding dropwise 20 mM sodium oleate in a prewarmed 5% solution of BSA in DPBS. The solution was sterile filtered and stored at −20°C. The solution of OA-BSA complexes will be referred to as “oleic acid” or “OA” in the following. Cells were treated with 400 μM OA in culture medium for 11-24 h.

### 2.5 Immunocytochemistry

#### 2.5.1 Immunostaining of hippocampal cultures

Hippocampal cells were fixed between day *in vitro* 12 and 21 in 4% paraformaldehyde in phosphate buffered saline for 20 minutes at room temperature. Cells were permeabilized and blocked with saponin-based buffer (SBB), containing 1% BSA and 0.1% saponin (Sigma) in PBS, for 30 minutes. The cells were incubated overnight at 4°C with primary antibodies in SBB: rabbit anti Myc (Myc; 1:1000; Abcam Cat# ab9106, RRID:AB_307014), rabbit anti Mover (Mover; 1:1000; Synaptic Systems Cat# 248 003, RRID:AB_10804285), guinea pig anti perilipin-2 (Perilipin-2; 1:1000; Progen Cat# GP40, RRID:AB_2895086), mouse anti glial fibrillary acidic protein (GFAP; 1:2000; Synaptic Systems Cat# 173 011, RRID:AB_2232308), guinea pig anti synaptophysin 1 (Synaptophysin; 1:1000; Synaptic Systems Cat# 101004, RRID:AB_1210382), chicken anti microtubule associated protein 2 (MAP2; 1:1000; Synaptic Systems Cat# 188 006, RRID:AB_2619881). After washing in PBS, cells were incubated 1 hour at room temperature in the dark with secondary antibodies and LD probes in SBB: goat anti chicken conjugated to Cy3 (1:500 – 1:1000; Jackson Cat# 703-166-155, RRID:AB_2340364), goat anti guinea pig conjugated to Cy2 (1:1000; Jackson Cat# 106-225-003, RRID:AB_2337427), donkey anti guinea pig conjugated to Cy3 (1:500 – 1:2000; Jackson Cat# 706-165-148, RRID:AB_2340460), goat anti mouse conjugated to Cy2 (1:1000 – 1:2000; Jackson Cat# 105-225-146, RRID:AB_2307343), donkey anti rabbit conjugated to Alexa 647^™^ (1:2000; Livetec Cat# A31573, RRID:AB_2536183), HCS LipidTox^™^ Red Neutral Lipid Stain (LipidTox, 1:1500; Invitrogen), BODIPY^™^ 493/503 (Bodipy, 10 μM; Molecular Probes). To label the nuclei, the cells were stained with 40≰6-diamidino-2-phenylindole (DAPI; Serva) in ddH2O for three minutes. Coverslips were mounted on glass microscope slides (Thermo scientific) in mowiol (Hoechst) with 2.5% 1,4-diazabicyclo[2.2.2]octane (Merck, DABCO). For STED microscopy Mover was labelled using the secondary antibody goat anti rabbit STAR 635P (1:100; Abberior, Cat# ST635P-1002-500UG, RRID:AB_2893229).

#### 2.5.2 Immunostaining of astrocyte cultures

Cortical astrocytes were fixed in 4% paraformaldehyde in phosphate buffered saline for 20 minutes at room temperature. Primary antibodies were incubated overnight at 4°C in 2% BSA + 10% Fetal Calf Serum (FCS) + 0.3% Triton-X-100: mouse anti GFAP (GFAP; 1:4000; Synaptic Systems Cat# 173 011, RRID:AB_2232308), rabbit anti Mover (Mover; 1:1000; Synaptic Systems Cat# 248 003, RRID:AB_10804285), ionized calcium adaptor molecule 1 (IBA1; 1:2000; Synaptic Systems Cat# 234 006, RRID:AB_2619949). Excess of primary antibodies were washed out with PBS. Secondary antibodies and LDs markers were incubated 1 hour at room temperature in dark: donkey anti mouse conjugated with Alexa Fluor^™^ 647 (1:1000; (Jackson ImmunoResearch Labs, Cat# 715-605-151, RRID:AB_2340863), donkey anti rabbit conjugated with Cy5 (1:500; Jackson ImmunoResearch Labs, Cat# 711-175-152, RRID:AB_2340607), HCS LipidToxTM Red Neutral Lipid Stain (LipidTox, 1:200; Invitrogen). For STED microscopy the protein Mover was stained with goat anti rabbit STAR 635P (1:250; Abberior, Cat# ST635P-1002-500UG, RRID:AB_2893229). Coverslips were mounted on glass microscope slides (Thermo scientific) in mowiol (Hoechst) with 2.5% 1,4-diazabicyclo[2.2.2]octane (DABCO; Merck).

### 2.6 Microscopy

#### 2.6.1 Epifluorescence microscopy

Images were acquired using a Zeiss Axio Imager.Z2 epifluorescence microscope equipped with the 16-bit 0RCA-flash4.0 V2 digital Cmos camera (Hamamatsu) and ZEN Blue software version 2.3. The Apotome.2 was used for optical sectioning of HEK cells.

#### 2.6.2 Super resolution STED microscopy

Stimulation emission depletion (STED) microscopy was performed using two different setups. A Leica DM18 microscope (Leica Microsystems) equipped with a titanium–sapphire Laser (MaiTai; Spectra-Physics, Darmstadt, Germany) was used for endogenous Mover imaging around LDs in primary hippocampal cultures. The setup was based on the setup used by Göttfert *et al*. (2013). Briefly, excitation beams at 568/20 nm (Bright Line HC, Semrock, Rochester, NY) or 650/13 nm (Bright Line HC, Semrock) were generated using bandpass filters. STED pulses at 775 nm were generated with a pulsed laser (Katana 08 HP, OneFive GmbH, Regensdorf, Switzerland). An Abberior Expert Line STED Instrument microscope using avalanche photodiodes (APD) detectors was used for transfected Mover in astrocyte cultures. STED images were acquired with UPLSAPO-HR 100x APO, 1.4 NA immersion objective in oil. The samples were excited with a 561 nm, 40 MHz pulsed excitation laser (Abberior Instruments) for LipidTox and a 640 nm, 40 MHz pulsed excitation laser (Abberior Instruments) for STAR 635P. A STED laser 775 nm (1.2W) pulsed laser from M PB Communications Inc., was used for STED acquisitions. Images were acquired with the Imspector software.

### 2.7 Statistic

The number and size of ring-shaped Mover structures were quantified using the multi point tool from FIJI (Schindelin *at al*. 2012). The selection of the astrocyte cells was based on GFAP-positive staining. As the immunofluorescence signal of Mover in epifluorescence images produced closed ring-shaped structures, those structures were referred to as “Mover rings”. A selection of Mover rings per region was gained by a blinded random approach, and the diameter of these selected rings were measured. The diameters were determined in microns using FIJIs straight tool. Mover rings were counted in 52-84 whole regions of 154.76 x 154.76 μm each on coverslips of four independent preparations (n=4). For quantification, results were grouped in untreated (−OA) and OA treated (+OA) cells. In GraphPad Prism 5, results were arranged in scatter plots showing the mean and the standard error of mean (SEM). For LD quantification, structures were counted as LDs if they were visible by differential interference contrast (DIC) and contained a positive LipidTox staining. ROIs were selected based on the GFP staining. LDs in Mover overexpressing group and control group were counted using the multi point tool from ImageJ while blinded for the condition. For data analysis, the two-tailed Wilcoxon-Mann-Whitney test for independent samples, p-value = 0.05, was performed in GraphPad Prism 5 and R Studio version 1.4.1717 and R language version 4.1.0.

## 3 Results

### 3.1 Mover formed ring-shaped structures in astrocytes

In primary hippocampal co-cultures, containing neurons and astrocytes, Mover co-localizes with the presynaptic marker protein synaptophysin (Kremer *et al*. 2007; Ahmed *et al*. 2013) (Figure 1a-e, arrowheads). However, we unexpectedly observed ring-shaped Mover staining outside MAP2-positive neurons (Figure 1a-e, arrow). These ring-shaped Mover structures were then found to be present in GFAP-positive astrocytes (Figure 1f-j). Extraneural Mover structures were remarkably larger than synaptic Mover puncta (Figure 1b-e arrowheads) and were not associated with the synaptic protein synaptophysin (Figure 1b-e arrow and g-j). These observations suggested that Mover is not only a presynaptic protein but is also associated with organelles in astrocytes. Because of their characteristic shape, we suspected that these ring-shaped structures might represent LDs. As the number and size of LDs can be increased by treating cells with excess fatty acids (Nakajima *et al*. 2019), we applied oleic acid in astrocytes to see if this affects the ring-shaped Mover structures in a similar manner.

**Figure 1:**
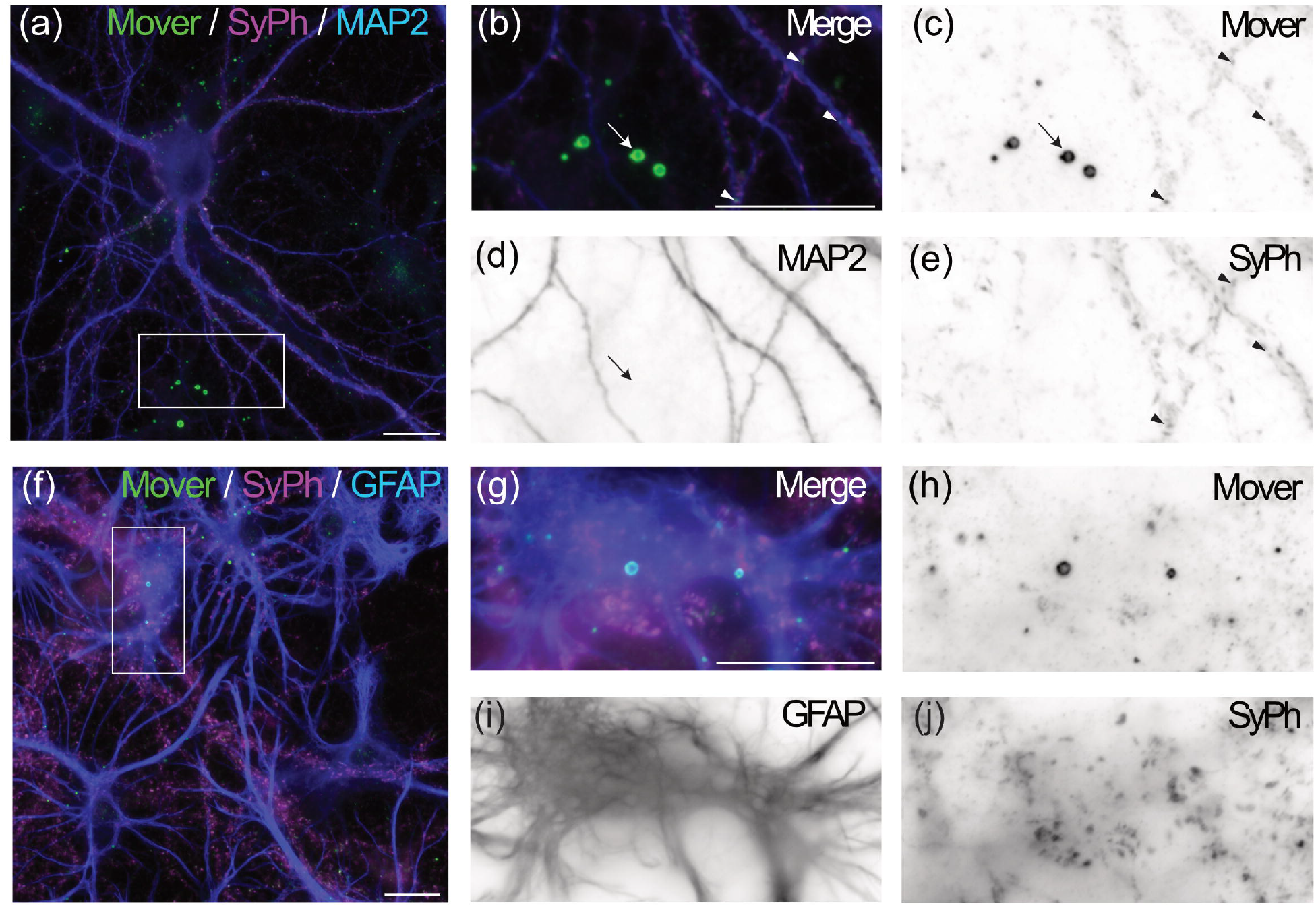
Mover immunofluorescence appears as ring-shaped structures in astrocytes. (a) Epifluorescence microscopy of a hippocampal neuron-astrocyte culture stained with MAP2 for neurons (blue), synaptophysin (magenta), and Mover (green), (b-e) Zoom on the inlet in (a), highlighting the ring-shaped Mover structures; merge (b) and single channel images (c-e). The arrow indicates the extraneuronal ring-shaped Mover structure; the arrowheads indicate colocalization of synaptic Mover with synaptophysin around MAP2 stained neurons. (f) Epifluorescence microscopy of hippocampal neuron-astrocyte culture; stained with GFAP for astrocytes (blue), synaptophysin (magenta), and Mover (green). (g-j) Zoom on the inlet in (f); merge (g) and single channel images (h-j). The ring-shaped Mover structure is located within the GFAP stained astrocytes. Scale bars = 20 μm.

In untreated astrocytes, ring-shaped Mover structures appeared small and low in numbers (Figure 2a-d), whereas application of OA rapidly increased their number and size (Figure 2eh). We quantified these differences and observed a significant higher number (Figure 2i) and increase in the diameter (Figure 2j) (number: p< 0.0001, Mann-Witney-U-test, two-tailed; diameter: p< 0.0001, Mann-Witney-U-test, two-tailed) of ring-shaped Mover structures in astrocytes after OA treatment): Untreated cells showed an average of 16 ± 0.73 ring-shaped Mover structures per region with an average diameter of 1.04 ± 0.03 μm (mean ± SEM), compared to 70 ± 3 ring-shaped Mover structures with an average diameter of 1.35 ± 0.03 μm (mean ± SEM) in OA treated cultures (Figure 2i,j). These experiments demonstrated striking similarities in the behavior of LDs and ring-shaped Mover structures that led us to further investigate their identity.

**Figure 2:**
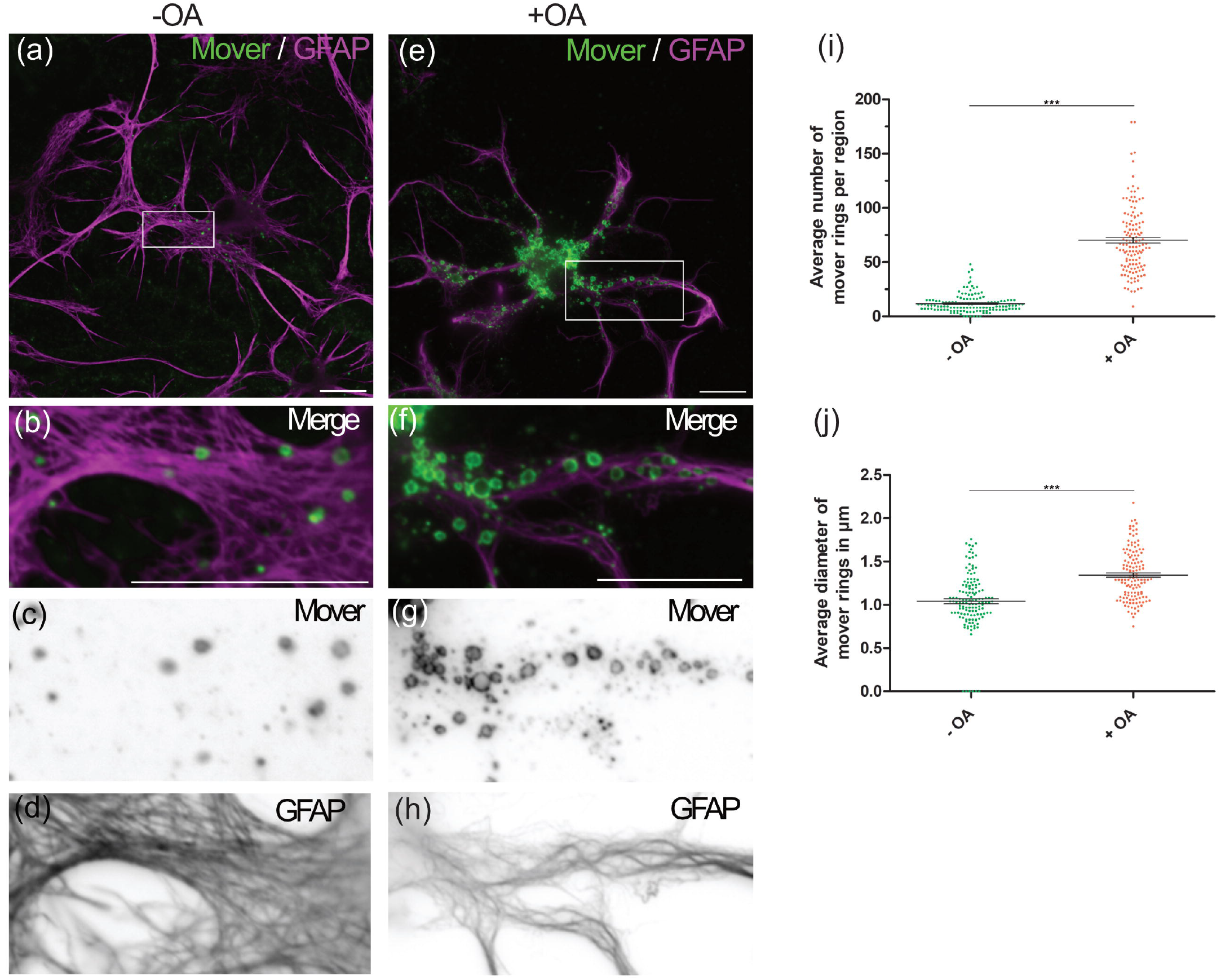
Oleic acid treatment increases the number and size of Mover-positive ring-shaped organelles. (a) Representative overview of untreated hippocampal astrocytes stained with GFAP (magenta) and Mover (green). (b-d) Zoom on the inlet in (a) show few small ring-shaped structures within the astrocyte; merge (b) and single channel images (c-d). (e) Representative overview of OA treated hippocampal astrocytes stained with GFAP (magenta) and Mover (green). (f-h) Zoom on the inlet in (e) show larger ring-shaped Mover structures within the astrocyte; merge (f) and single channel images (g-h). (i) Representation of the number of ring-shaped Mover structures per region in absence (−OA) or presence (+OA) of OA treatment. (j) Representation of the diameter of ring-shaped Mover structures per region in absence (−OA) or presence (+OA) of OA treatment. Bars show mean ± SEM. Statistical evaluation was done using a two-tailed Wilcoxon-Mann-Whitney test for 52-84 regions in 3-4 different preparations (n=3-4). Asterisks represent significant differences with *** = p ≤⍰0.001. Scale bars = 20 μm.

### 3.2 Mover is associated to LDs

To confirm the presence of Mover around LDs we stained OA treated cells with different LD markers. The LD associated protein perilipin-2 (Heid *et al*. 1998) was selected as specific staining of the LD surface. Mover was associated with both perilipin positive (Figure 3, arrows) and perilipin negative structures (Figure 3, arrowheads). Thus, one pool of Mover is associated with LDs while another pool of Mover may be associated with different organelles or with perilipin negative LDs. To test this, we labelled the LDs neutral lipid core using the fluorescent dyes Bodipy and LipidTox.

**Figure 3:**
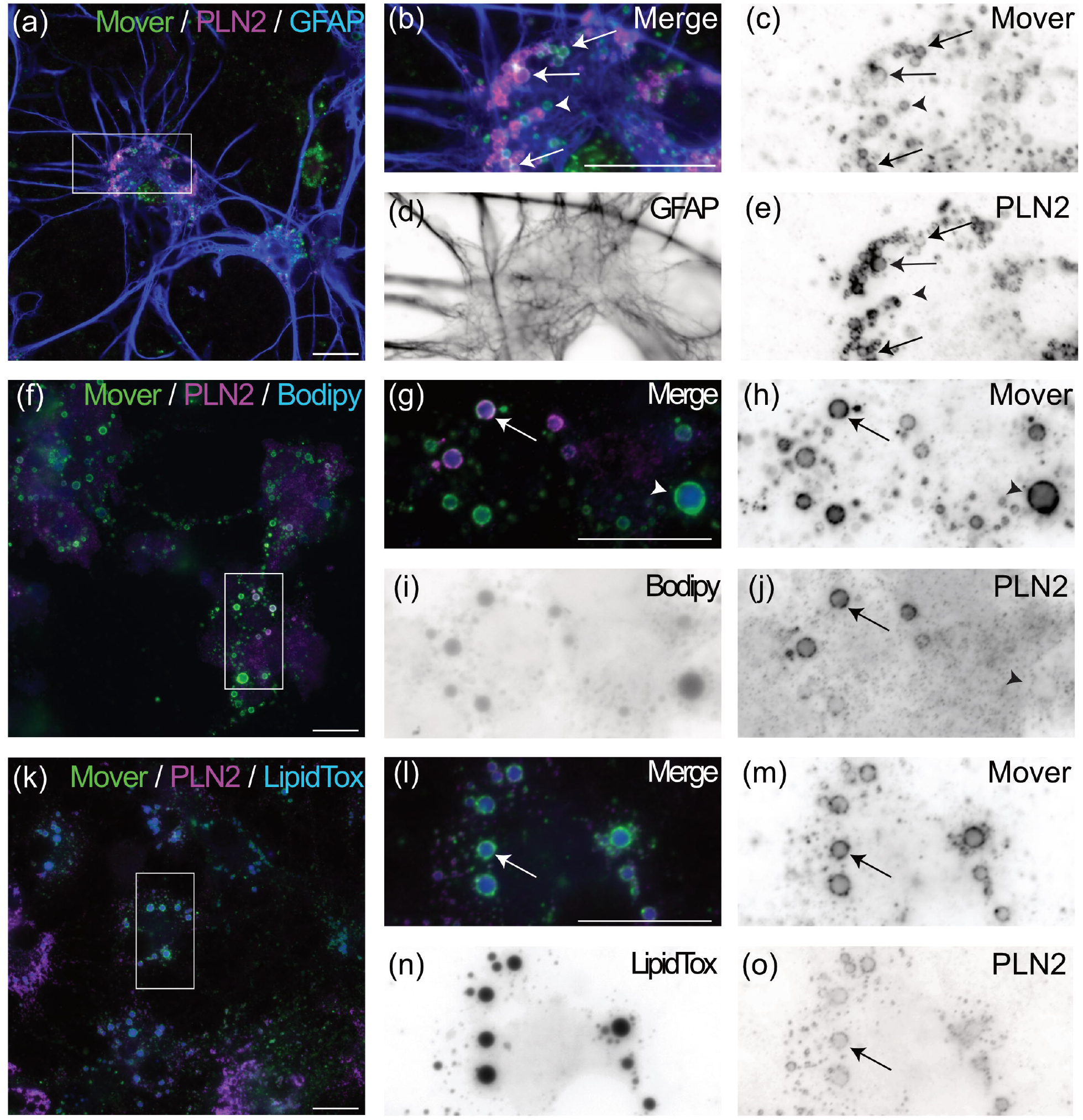
Colocalization of Mover with LD markers Bodipy, LipidTox and Perilipin-2. (a) Hippocampal astrocytes stained for Mover (green), GFAP (blue), and perilipin-2 (magenta). (b-e) Zoom on the inlet in (a) showing partial overlap of Mover with perilipin-2. Merge (b) and single channel images (c-e). (f) Hippocampal astrocytes stained for Mover (green), Bodipy (blue), and perilipin-2 (magenta), (f-j) Zoom on the inlet in (f) showing overlap of Mover with LD marker Bodipy. Merge (g) and single channel images (h-j). (k) Hippocampal astrocytes stained for Mover (green), LipidTox (blue), and perilipin-2 (magenta). (l-o) Zoom on the inlet in (k) showing overlap of Mover with LD marker LipidTox. Merge (l) and single channel images (m-o). Arrows indicate overlap of Mover and perilipin-2, while arrowheads indicate Mover positive, but perilipin-2 negative structures. Scale bars = 20 μm.

Virtually all ring-shaped Mover structures surrounded accumulation of neutral lipids, as verified by both Bodipy (Figure 3f-j) and LipidTox stainings (Figure 3k-o). These results indicate that Mover localizes to the surface of astrocytic LDs and that some of them are not associated with perilipin-2.

STED microscopy further supports the observation that ring-shaped Mover is associated to LDs (Figures 4 and 5). Endogenous Mover does not appear as closed ring around OA induced LipidTox-positive LDs, but as a broadly distributed punctate staining around the LD surface (Figure 4a-d), similar to previously reported results for the LD proteins perilipin-2 and perilipin-5 (Gemmink *et al*. 2018). Furthermore, we occasionally observed that Mover formed local clusters on the LD surface (Figure 4e-g) or appears at the contact site between LDs (Figure 4h). These result raises the possibility that Mover is arranged in nanodomains on their surface. We then stained for Mover immunofluorescence in cells that expressed monomeric GFP (mGFP) as a control (Figure 5a) or the recombinant mGFP-Mover (tagged with mGFP at the N-terminus) (Figure 5b). As expected, the antibody revealed punctate staining for endogenous Mover in mGFP expressing cells (Figure 5a). In contrast, the immunostaining covered the entire LD surface in mGFP-Mover expressing cells (Figure 5b). This suggests that there is additional capacity for Mover recruitment on the LD surface in control cells, which recombinant Mover can occupy. Moreover, in mGFP-Mover overexpressing cells, the LDs appeared to be easier to detect than in control cells, even without OA (Figure 5). This led us to hypothesize that overexpressing Mover affects LDs.

**Figure 4:**
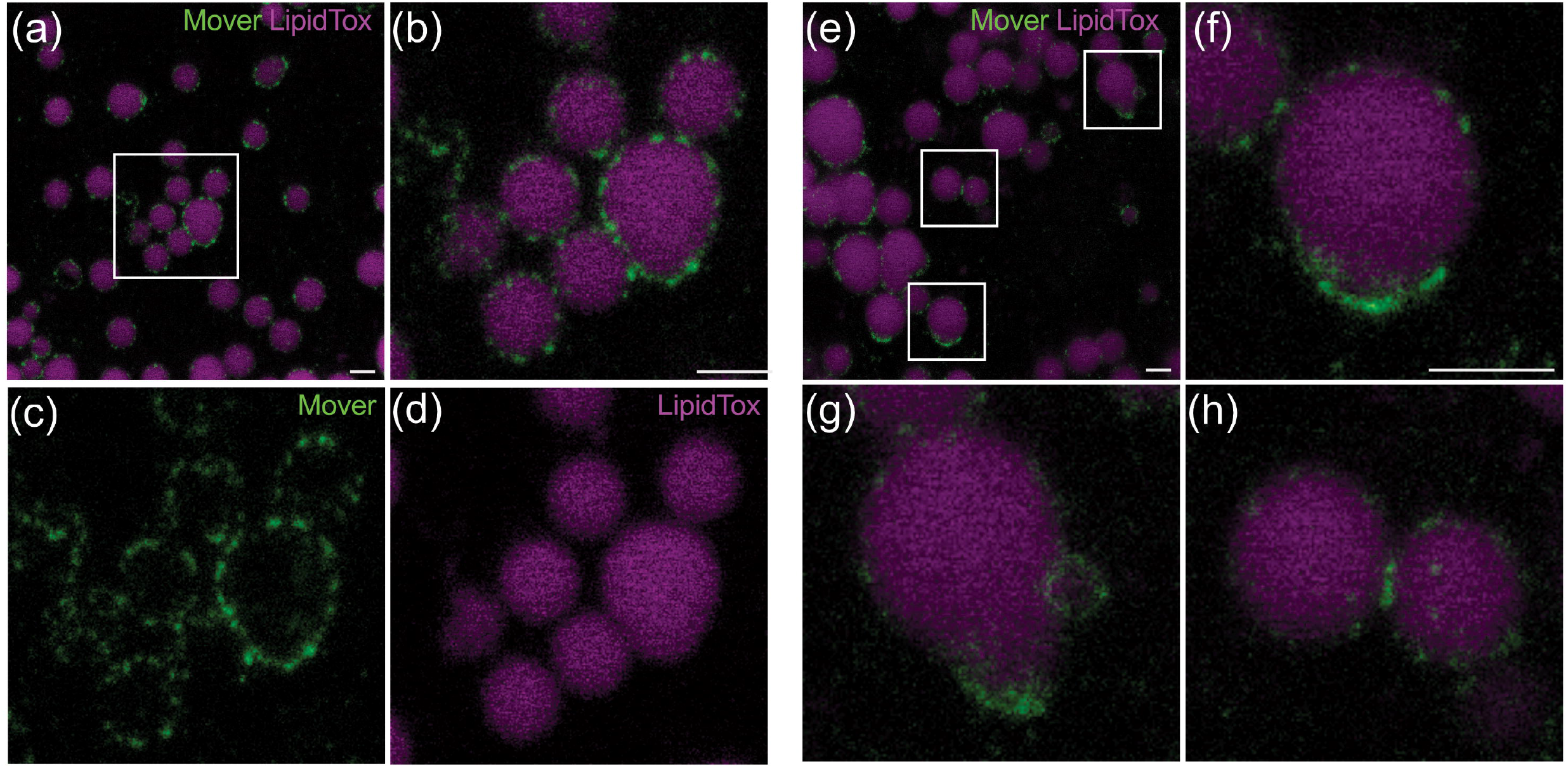
STED-microscopy reveals association of endogenous Mover with LDs. (a) OA induced LDs from hippocampal cultures stained with LipidTox (magenta) and for Mover (green) imaged by STED microscopy. (b-d) Zoom on the inlet shown in (a). Merge (b) and single channel images (c-d) show the distribution of endogenous Mover around the LDs and the neutral lipid staining inside. (e) OA induced LDs from hippocampal cultures stained with LipidTox (magenta) and Mover (green) imaged by STED microscopy, (f-h) Zoom of three regions of interest shown in (e) indicating the distribution of Mover in nanodomains around LDs. Scale bars = 1 μm.

**Figure 5:**
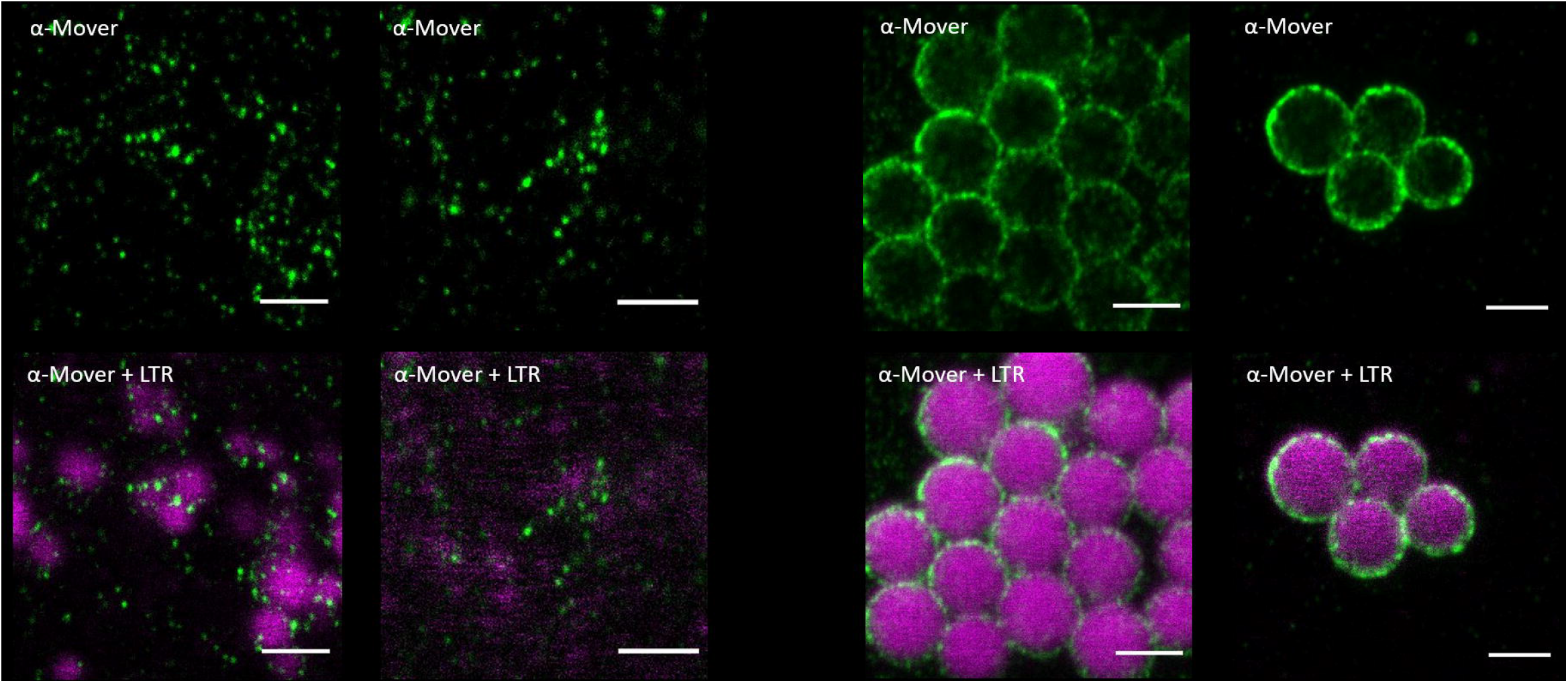
STED microscopy shows targeting of recombinant Mover to the LD surface in astrocyte cultures. a) Distribution of Mover immunofluorescence (green) around LipidTox (LTR) stained LDs (magenta) in mGFP transfected control astrocytes in presence (+OA) of oleic acid treatment compared to untreated (−OA) cells, b) Distribution of Mover immunofluorescence (green) around LipidTox (LTR) stained LDs (magenta) in recombinant-Mover overexpressing astrocytes in presence (+0A) or in absence (−OA) of oleic acid treatment. Mover overexpression was sufficient to cause a marked increase in the visibility and contrast of LDs compared to untreated mGFP transfected control, where LDs appear shapeless. Scale bars = 1 μm.

### 3.3 Effect of Mover overexpression on LD formation in astrocytes

In order to perform a quantitative gain of function assay, we generated a new plasmid encoding untagged Mover together with an IRES driven membrane targeted GFP as reporter. We used the GFP signal to delineate the borders of transfected cells. A plasmid encoding only a membrane-targeted GFP (GFP-F) was used as a control.

We then expressed these constructs in astrocyte cultures and quantified the number of LDs within transfected cells (Figure 6a). To apply stringent criteria for quantification we only regarded structures as LDs when they were both LipidTox-positive and visible in DIC (Figure 6b). We observed significantly more LDs in the cells overexpressing Mover compared to control cells expressing GFP-F (Figure 6a and b) (Mover overexpressing cells: 132 ± 75 (mean ± SEM); control cells: 26 ± 23 (mean ± SEM); (p-value < 0.0001, Mann-Witney-U-test, twotailed)) (Figure 6c). This result explains why the LDs were easier to detect in non-OA treated cells overexpressing Mover (Figure 5b) and demonstrates that Mover is sufficient to increase the number of LDs in astrocytes.

**Figure 6:**
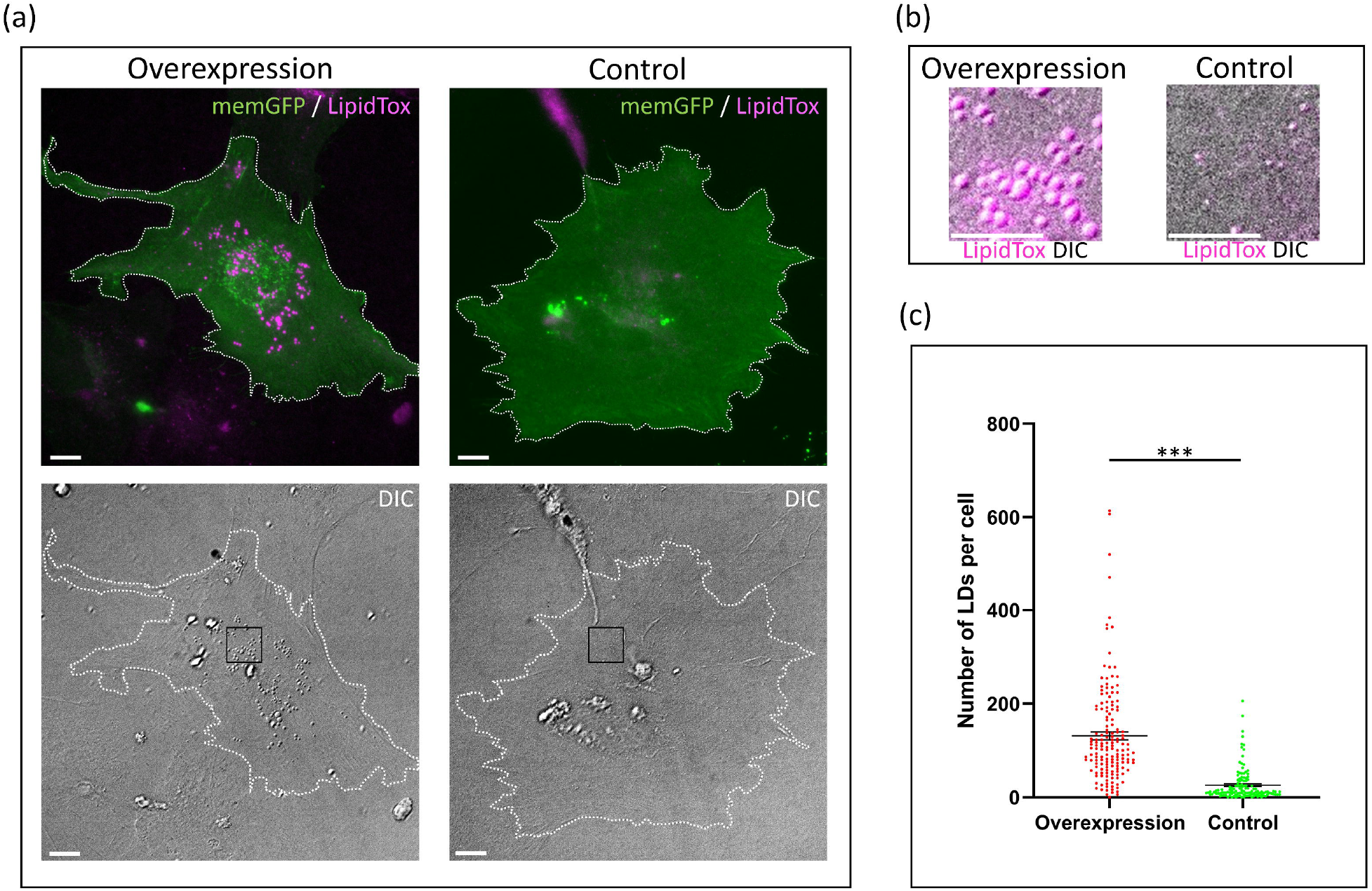
Mover overexpression induces accumulation of LDs in astrocytes. a) Astrocytes were transfected either with a recombinant Mover-GFP construct (overexpression) or with a GFP control (control). LDs were stained with LipidTox (magenta). Astrocytes were delineated (white dashed line) using the membrane targeted GFP signal (memGFP) (green); bottom panels show the respective DIC channel. b) Zoom on the inlets in DIC channel in (a) reveals LD accumulation after Mover overexpression, c) Quantification of the LD number per astrocyte in overexpression and control condition reveals a marked increase of LDs after Mover overexpression. Bars represent mean ± SEM. Statistical evaluation was done using a two-tailed Wilcoxon-Mann-Whitney test. Asterisks represent significant differences with *** = p ≤⍰0.001. Scale bars = 10 μm.

### 3.4 Identification of Mover sequences necessary for LD association

To determine Mover sequences necessary for LD association, we transfected HEK293 cells with recombinant Mover constructs. We investigated the ability of each recombinant Mover construct to associate with LDs after OA treatment. In addition to the N-terminal tagged mGFP-Mover (Figure 7a-d), the two C-terminal tagged Mover constructs Mover-mGFP (tagged with monomeric GFP) (Figure 7e-h) and Mover-eGFP (tagged with conventional eGFP) (Figure 7i-l) co-localized around LDs. To exclude a contribution of the GFP tag to the LD association, we also tested a C-terminal myc-tagged Mover construct that showed association to LDs after staining for Mover (Figure 7m-p). These results suggest that Mover does not need a free N-terminal or C-terminal domain to interact with LDs. Likewise, a truncated Mover construct missing the first 51 amino acids (mGFP-52-266) still associated with LDs (Figure 7q-t), which indicates that the N-terminal 51 amino acids are not required for LD binding. We then explored if phosphorylation is necessary for LD association and expressed a phosphorylation-deficient Mover construct with mutations at five predicted phosphorylation sites (Mover-5p-eGFP, including the following mutations: T13A, S14A, T64A, S221A, Y257F). This phosphorylation-deficient mutant did not lose its ability to associated with LDs either (Figure 7u-x), suggesting that Mover does not require phosphorylation at these five distinct sites to bind to LDs.

**Figure 7:**
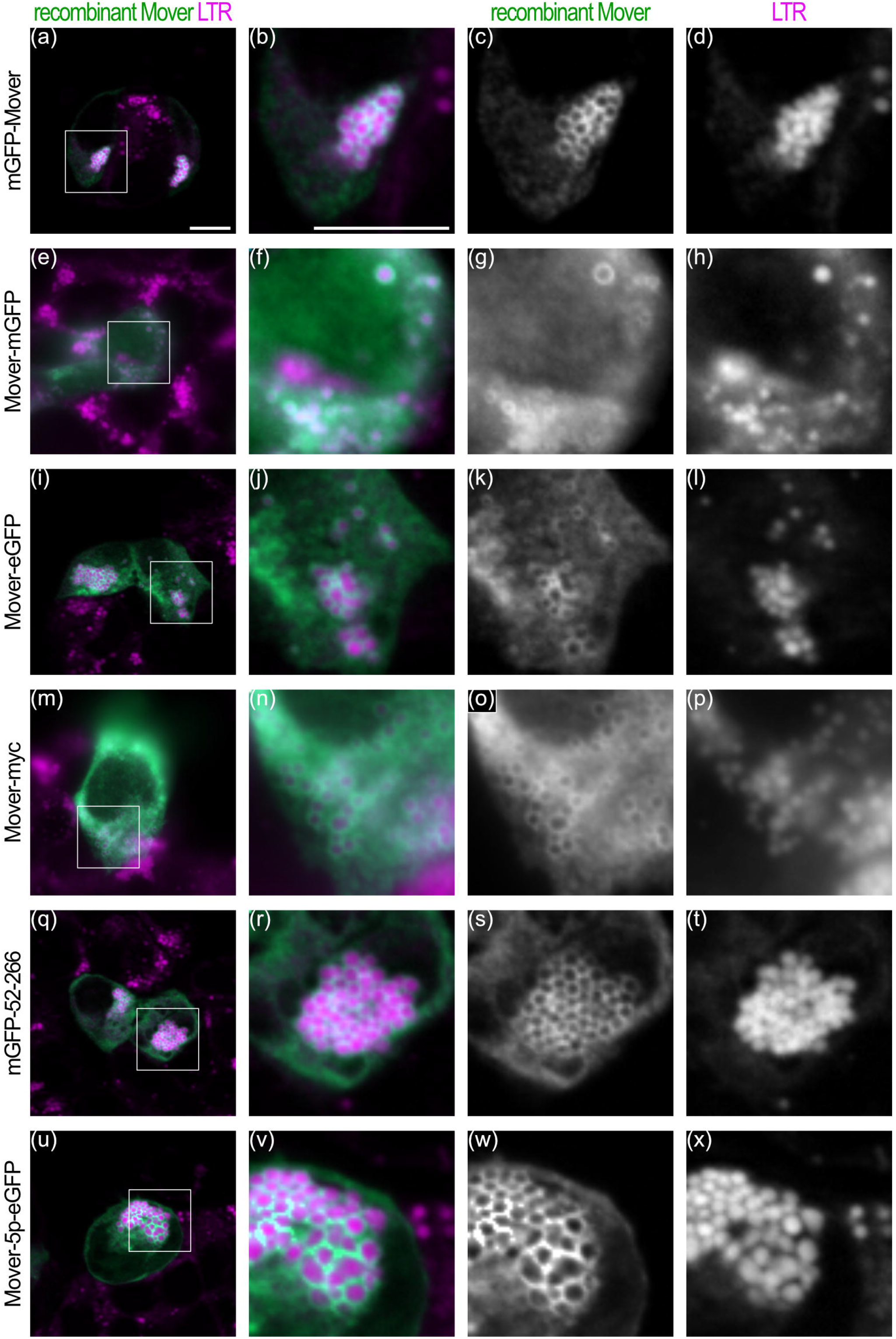
Different recombinant Mover constructs associate to LDs in HEK293-cells. OA treated HEK cells were transfected with mGFP-Mover (a-d), Mover-mGFP (e-h), Mover-eGFP (i-l), Mover-myc (m-p), mGFP-52-266 (q-t), and Mover-5p-eGFP (u-x). Recombinant Mover signal (green) from the respective GFP tag, except for Mover-myc, where Mover antibodies were used, and LipidTox (LTR) stained LDs (magenta) show strong association of different constructs to LDs. First panel per row on the left gives an overview, the other panels zoom on the indicated inlet with a merge and the respective single channel images. Scale bars = 10 μm.

To further explore regions required for LD association, we tested three possibilities arising from our previous work. First, introducing four point mutations (F206R, K207E, K215E, K219E) abolishes calmodulin binding of Mover (Akula *et al*. 2019). A construct harboring these mutations (Mover-4cam-eGFP) failed to associate with LDs (Figure 8a-d). Interestingly, a single point mutation (F206R) was sufficient to abolish LD association, both in full length (Figure 8e-h) and in the truncated construct mGFP-51-266 (Figure 8i-l). These results indicate that F206 is essential for LD association. Second, *in silico* analysis predicts a conserved domain, called hSac2 domain, spanning amino acids 53-163 of Mover. A construct encoding only the hSac2 domain did not bind to LDs (Figure 8m-p). Third, database entries suggest that at the cDNA level a Mover isoform exists that lacks exon 2, which encodes amino acids 94-150 and contains a large part of the hSac2 domain. A construct lacking exon 2, and thereby disrupting the hSac2 domain, failed to bind to LDs (Figure 8q-t). Together these data indicate that in addition to F206 the amino acids 94-150 are required for LD association. Moreover, they show that the hSac2 domain is required but not sufficient for LD binding.

**Figure 8:**
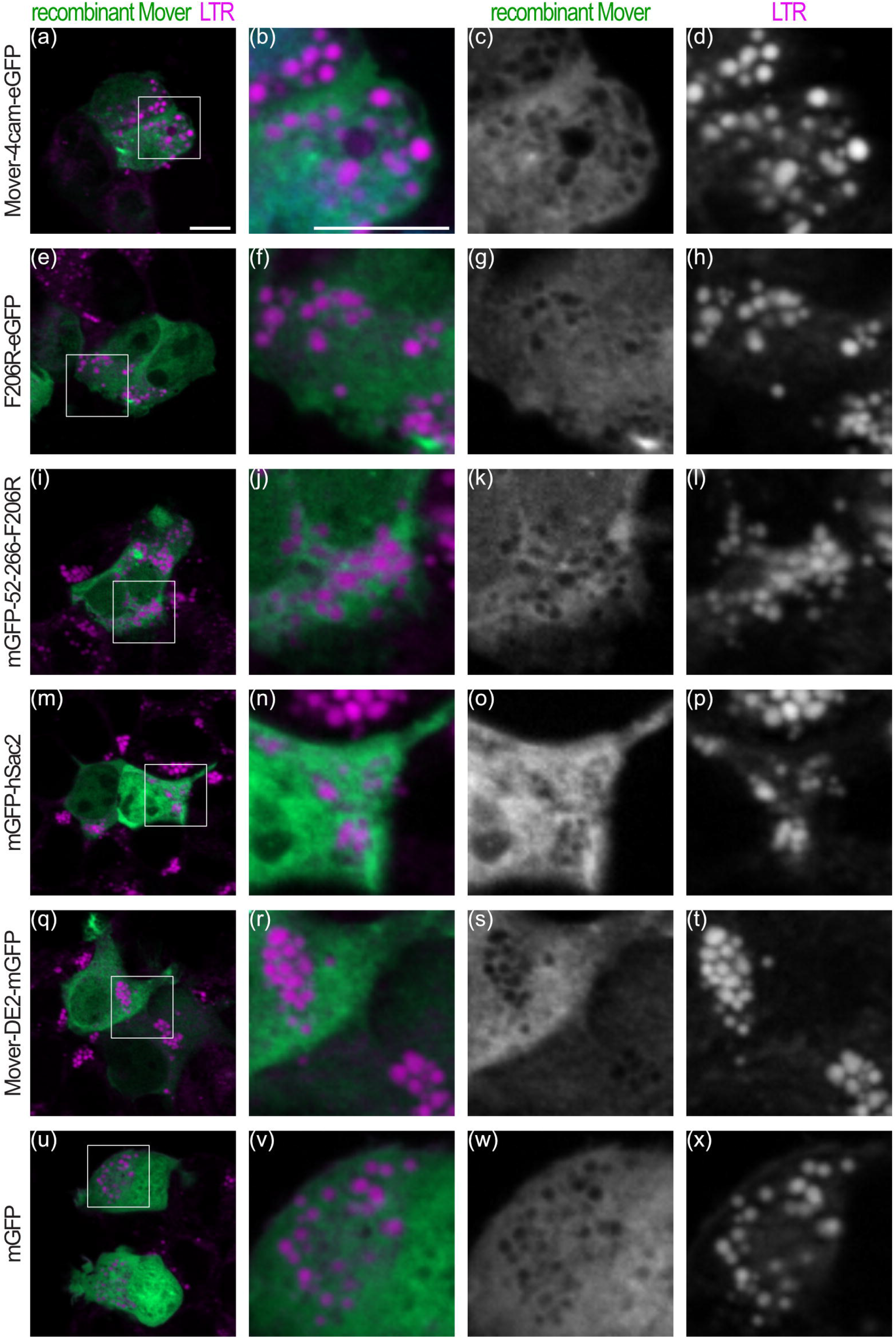
Extended regions of Mover are required for targeting to LDs in HEK293-cells. OA treated HEK cells were transfected with Mover-4cam-eGFP (a-d), F206R-eGFP (e-h), mGFP-52-266-F206R (i-l), mGFP-hSac2 (m-p), Mover-DE2-mGFP (q-t), and mGFP (u-x). Recombinant Mover signal (green) from the respective GFP tag and LipidTox (LTR) stained LDs (magenta) reveal no association of the probed constructs with LDs, similar to GFP control. First panel per row on the left gives an overview, the other panels zoom on the indicated inlet with a merge and the respective single channel images. Scale bars = 10 μm.

All deletion constructs that failed to bind to LDs were distributed in a way similar to soluble mGFP (Figure 8u-x).

## 4 Discussion

Astrocytic LDs protect surrounding neurons against lipid stress. The biogenesis and function of LDs relies on the composition of their protein coat. In this study, we identified the presynaptic protein Mover as a novel component of astrocytic LDs using different LD markers. We showed that the association with LDs is disturbed by large deletions in the exon 2 region and point mutations in the calmodulin binding region of Mover. Furthermore, overexpression of Mover induces LD accumulation in astrocytes, highlighting Mover as a candidate regulator of astrocytic LDs.

### 4-1 Mover is a novel component of astrocytic LDs

Several lines of evidence indicate that Mover is a protein of the LD surface in astrocytes. First, using epifluorescence microscopy, we found endogenous Mover surrounding LDs identified by two distinct neutral lipid stains LipidTox and Bodipy. Similarly, recombinant Mover retained the ability to bind to LipidTox-positive LDs, both in astrocytes and HEK cells. To strengthen our observation that Mover is part of the LD coat, we labeled perilipin-2, a known part of the LD surface, and saw that it colocalized well with Mover. Second, after OA treatment, Mover behaves as expected for a LD associated protein, in that the number and size of the ring-shaped structures formed by Mover increased. Third, super-resolution STED microcopy confirmed the presence of endogenous and recombinant Mover on the LD surface. The punctate STED signals found across the LD surface is reminiscent of the STED signal of perilipin-2 (Gemmink *et al*. 2018). However, while endogenous Mover was organized in nano-clusters on the LDs, perilipin-2 signal appeared to be more homogenously distributed.

Interestingly, we observed LDs and even entire cells with negligible perilipin-2 signals but prominent Mover-positive LDs. Thus, Mover may be a more widespread marker for astrocytic LDs compared to perilipin-2. Moreover, we conclude from this observation that not all astrocytic LDs exhibit the same surface protein coating. The differences we observed are consistent with results from CHO K2 cells, Huh7 cells, in mouse liver, and in mouse brown adipose tissue, where LDs were separated by size and distinct surface protein compositions (Zhang *et al*. 2016). Coat identity could depend on the size of the LD, the time elapsed after their formation, or their interaction with a distinct organelle in the cell. Furthermore, the brain-specific LD protein GRAF1a shows similarities with Mover in this respect: recombinant GRAF1a surrounded most astrocytic LDs after OA treatment, and a subset of GRAF1a-positive LDs were devoid of perilipin-2 or perilipin-3 (Lucken-Ardjomande Häsler *et al*. 2014). In other aspects, Mover differs from GRAF1a: STED microscopy revealed GRAF1a is mainly found at LD contact sites but also forms long vesicular structures. In contrast, we found that Mover forms nanodomains across the entire LD surface. In addition, we demonstrated that endogenous Mover is found around astrocytic LDs, without need of overexpression. From these results we conclude that, while Mover shows characteristics similar to GRAF1a, it localizes to different substructures on the LD, suggesting that Mover is involved in different mechanisms.

Synaptic vesicles and LDs are highly specialized organelles that are distinct in their structure, function, and biogenesis. One major difference is their membrane structure: while synaptic vesicles have a phospholipid bilayer membrane, LDs exhibit a phospholipid monolayer membrane. Thus, the presence of Mover around both synaptic vesicles and LDs is intriguing. The dual association of Mover both structures is reminiscent of Annexin VI, which was first identified as interaction partner of the synaptic vesicle protein synapsin 1 (Inui *et al*. 1994), and more recently has been identified as a LD binding protein involved in hepatocytic LD formation (Cairns *et al*. 2017). Likewise, α-synuclein, initially found to regulate synapse maintenance (Murphy *et al*. 2000), has also been shown to modulate the triacylglyceride turnover in brain LDs (Cole *et al*. 2002). Parkinson’s disease related α-synuclein variants partially lost their affinity for LDs or their ability to regulate triacylglyceride metabolism (Cole *et al*. 2002), likely contributing to the disease. Interestingly, both α-synuclein and Mover are proteins that have uniquely evolved in vertebrates, suggesting vertebrate-specific functions.

### 4-2 Binding of Mover around LDs involves its predicted Calmodulin domain

Several modes of binding to LDs have been described: some proteins containing hairpin structures translocate from the ER to LDs during LD budding (Kory *et al*. 2017); GRAF1a binds to LDs via its membrane binding BAR domain (Lucken-Ardjomande Hasler *et al*. 2014); perilipins associate with existing LDs through n-terminal 11mer repeats that generate amphipathic helices (Rowe *et al*. 2016). Mover was not detected at the ER and does not contain a BAR domain. However, Mover contains a domain with structural similarity to pleckstrin-homology domains, called hSac2 domain, spanning amino acids 53-163 and covering the entire exon 2 (from amino acids 93-151). In the protein sac2, the hSac2 domain lacks phospholipid binding capacity, but supports subcellular targeting to early endosomes through dimerization (Hsu *et al*. 2015). In line with the observed lack of phospholipid binding, our results indicate that the hSac2 domain is not sufficient for the binding of Mover on the LD surface. On the other hand, deletion of exon 2, and thus removal of the hSac2 domain, prevented targeting of Mover to LDs in our study. This indicates that the hSac2 domain, while not sufficient, my contribute to the binding of Mover to LDs.

Strikingly, a single mutation exchanging the phenylalanine F206 for an arginine abolished the binding of Mover to LDs. This phenylalanine is localized in the calmodulin-binding sequence of Mover. This sequence is predicted to form an amphipathic helix. Thus, Mover may use a binding mode reminiscent of the mode used by perilipins, where amphipathic helices are the major binding sites. Intriguingly, phenylalanine F206 is also essential for both dimerization and targeting of Mover to the presynapse (Akula *et al*. 2019). Therefore, this short region of Mover is apparently involved in multiple functions, including calmodulin (CaM) binding, dimerization, targeting to presynaptic terminals in neurons, and targeting to LDs in astrocytes. A simple explanation for the importance of F206 would be that its hydrophobicity contributes to lipid binding or insertion within the phospholipid layer. Alternatively, F206 may be important for the overall folding of Mover.

### 4-3 Mover overexpression induces LDs accumulation in astrocytes

We found that Mover overexpression induces accumulation of LDs in astrocytes. Under normal conditions, LD number in cells is low, except for special fat storage cells such as adipose tissue. However, under stress, cells might accumulate LDs as a response and potential protective mechanism (Smolič *et al*. 2021). How Mover leads to the accumulation of LDs is not clear. The biogenesis of LDs comprises at least three steps: Triacylglycerides are synthesized in the ER and accumulate between the two membrane leaflets, leading to the budding of the ER as nascent LDs. Immature LDs then grow by additional triacylglyceride synthesis at the LD. Accordingly, the surface phospholipid layer needs to expand too (Wilfing *et al*. 2014; Olzmann & Carvalho 2019). Finally, LDs can grow by fusion with other LDs or by substrate exchange with other organelles, with which LDs form highly dynamic contacts (Valm *et al*. 2017). Although we did not observe any evidence of Mover localization to any organelle other than LDs, we cannot exclude that small fractions of Mover act at the ER. If Mover acts at a stage downstream of LD budding it may increase the size of existing LDs in a way that previously existing LDs become detectable after Mover overexpression. Mechanistically, this effect could involve increased triacylglyceride synthesis, reduced lipolysis, or increased LD fusion.

A hint to the pathways by which Mover may increase the number of detectable LDs comes from the fact that Mover may activate the transcription factor NF-κB (Liu *et al*. 2019; Lau *et al*. 2020). Pharmacological inhibition of NF-κB decreases lipid accumulation in HepG2 cells treated with high glucose (Daniel *et al*. 2021). This suggests that the activation of NF-κB is essential for the formation of LDs in hepatocytes. Thus, Mover may boost LD formation or growth by activating NF-κB signaling.

### 4-4 Mover might modulate neuroinflammation

The effect of Mover on astrocytic LDs discovered here and its reported effects on NF-κB signaling provide an interesting link to neuroinflammation. Impaired LD formation and function has been implicated in a large number of neurological conditions, spanning neurodegenerative disorders such as Alzheimer’s disease, Huntington’s disease, Parkinson’s disease, and Hereditary Spastic Paraplegia, as well as neuroinflammatory conditions (Farmer *et al*. 2020).

The protein FABP7 protects astrocytes from ROS toxicity by promoting LD formation (Islam *et al*. 2019). This is especially important, because astrocytes also take up toxic lipid species released from neurons to protect them from lipotoxicity (Ioannou *et al*. 2019). However, FABP7 overexpression in astrocytes also activates the NF-κB pathway and leads to a pro-inflammatory response that subsequently impairs motor neuronal survival *in vitro* (Killoy *et al*. 2020).

Induction of the transcription factors NFAT and NF-κB in astrocytes is a hallmark of both neuroinflammation and neurodegeneration (Fernandez *et al*. 2007; Sompol and Norris, 2018; Kraner & Norris 2018). Remarkably, Mover may activate both of these transcription factors: we found in a previous study that overexpression of Mover enhances the translocation of NFAT from the cytosol to the nucleus. (Akula *et al*. 2019). Nuclear translocation of NFAT is part of the NFAT activation pathway that results in the release of cytokines and pro-inflammatory signals. In addition, two studies have identified Mover (called TPRG1L in these studies) as being involved in the activation of the NF-κB pathway (Liu *et al*. 2019; Lau *et al*. 2020), which regulates the expression of pro-inflammatory genes such as cytokines and chemokines. TERC, the RNA component of telomerase, was shown to promote an inflammatory response. The authors suggested that TERC activates a set of genes, including Mover, which then stimulate the NF-κB pathway and thus elicit an inflammatory response (Liu *et al*. 2019). Accordingly, the expression levels of Mover were upregulated in patients suffering from inflammatory disorders such as multiple sclerosis or diabetes (Liu *et al*. 2019). Another study reported that the expression of Mover is upregulated in fibroblasts after infection with RNA1.2, the long-non-coding-RNA expressed by human cytomegalovirus. Interleukin (IL)-6 release was lower in Mover knock-down fibroblasts infected with RNA1.2, suggesting that Mover upregulation activates the NF-κB pathway (Lau *et al*. 2020). This highlights a possible implication of Mover in neuroinflammatory processes.

Inflammation with an increase of pro-inflammatory cytokines such as IL-6 or IL-8 and a dysregulation of the NF-κB pathway are prevalent in Schizophrenia (Murphy *et al*. 2021). In addition, Schizophrenia is characterized by lipid dysregulation (Maas *et al*. 2020). Strikingly, a proteomic screen of the anterior cingulate cortex, showed a 2.4 fold increase in expression of Mover (listed as novel protein RP11-46F15.3) in schizophrenic patients compared to healthy individuals (Clark *et al*. 2006). Together, the activation of the NF-κB pathway by Mover (Lau *et al*. 2020) and the increased levels of Mover in anterior cingulate cortex samples from schizophrenic patients (Clark *et al*. 2006) suggest an involvement of Mover in the development or progression of the disease, which could be related to the role of Mover at the synapse, or to its effect on LDs, or both.

The activity of both NFAT and NF-κB are modulated by Ca^2+^ signaling: Ca^2+^ activates CaM, which in turn activates the CaM-dependent phosphatase calcineurin. Activated calcineurin dephosphorylates NFAT, triggering its translocation to the nucleus. Calcineurin also indirectly activates NF-κB by acting on targets involved in NF-κB regulation (Sompol & Norris 2018). Mover binds CaM and stimulates the CaM/calcineurin pathway, actively enhancing NFAT1 translocation (Akula *et al*. 2019). Therefore, by activating CaM/calcineurin signaling, Mover could stimulate both NFAT and NF-κB, and thus lead to an increase of LDs in astrocytes. It will be interesting to see in future studies whether the increase in Mover expression, which occurs in schizophrenia, leads to an increase in astrocytic LDs and whether this reflects disorder-induced dysregulation or rather a protective role of Mover.

## 5 Conclusion

In summary our results demonstrate a novel localization for the presynaptic protein Mover on the surface of astrocytic LDs. Overexpression of Mover increased the number of LDs, suggesting that Mover affects LD biogenesis. This introduces Mover as a novel component of astrocytic LDs and a candidate regulator of the mechanisms driving accumulation of LDs in astrocytes.

## Acknowledgements

We thank Katrin Willig and Jakob Neef for their helpful support at their STED setups; and we thank Irmgard Weiß, Nina Dankenbrink-Werder and Lisa-Marie Hartmund for excellent technical assistance.

This study was supported by the DFG Center for Nanoscale Microscopy and Molecular Physiology of the Brain (CNMPB, B1-1 to TD).

## DATA AVAILABILITY STATEMENT

Data available on request from the authors

